# Reward evokes value-independent arousal and value-dependent reinforcement processing as dissociable physiological consequences

**DOI:** 10.64898/2026.08.17.745141

**Authors:** Yusuke Nakashima, Yuka Sasaki, Takeo Watanabe

**Affiliations:** Department of Cognitive and Psychological Sciences, Brown University, Providence, RI 02912

## Abstract

Reward is traditionally thought to drive value-dependent reinforcement processing, whereas recent work suggests reward may instead raise arousal that mimics reinforcement. Because reinforcement and arousal depend on partly distinct neuromodulatory systems yet are co-triggered by reward, they have been widely treated as closely linked, though rarely tested directly, leaving reward’s physiological consequences difficult to interpret. We manipulated the value of a primary reward by varying thirst while independently inducing arousal with sound, and measured pupil diameter and electroencephalographic activity. High-and low-value rewards produced comparable pupil dilation, indicating value-independent arousal. In contrast, only high-value reward increased alpha-band power, progressing from anterior to posterior regions, whereas low-value reward and sound produced similar alpha suppression without regional progression. Thus, reward evokes two dissociable consequences: value-independent arousal and value-dependent reinforcement. Rather than a single coupled process, these arise through separable mechanisms, providing a physiological framework for distinguishing each mechanism’s contribution to reward processing.

## Introduction

Reward shapes behavior through at least two processes. Traditionally, reward is thought to drive value-dependent reinforcement processing, in which dopaminergic systems represent the value of outcomes and use it to guide future behavior (Schultz et al., 1997; Schultz, 2016). More recent work, however, has emphasized that arousal itself can modulate cortical processing and facilitate learning (Mather & Sutherland, 2011; Nassar et al., 2012; Glennon et al., 2019). Reward reliably increases arousal (Sara, 2009), raising the possibility that some processing attributed to reinforcement instead reflects its arousing rather than reinforcement properties. Because a single rewarding event can engage both processes, it has been difficult to determine how each contributes to the physiological consequences of reward.

Reinforcement and arousal depend on partly distinct neuromodulatory systems. Dopaminergic neurons primarily encode reward value (Schultz et al., 1997; Schultz, 2016), whereas the locus coeruleus-norepinephrine (LC-NE) system plays a central causal role in regulating arousal and wakefulness (Berridge & Waterhouse, 2003; Carter et al., 2010; Berridge, 2012). Despite this separation, an influential body of work has suggested that the two are closely linked. Dopaminergic neurons encode not only reward value but also motivational salience and alerting signals (Matsumoto & Hikosaka, 2009; Bromberg-Martin et al., 2010), and the LC-NE system is itself engaged by reward-related events and modulates cortical processing through gain control (Aston-Jones & Cohen, 2005; Sara, 2009; Breton-Provencher et al., 2022). Consistent with this view, physiological arousal accompanies reward anticipation and delivery (Critchley et al., 2001), and arousal has been proposed to bias cortical processing toward reward (Mather & Sutherland, 2011). Together, these observations have led reinforcement and arousal to be widely treated as closely linked.

This assumed link, however, has rarely been tested directly, and there is a basic reason for caution: reward is intrinsically arousing. A rewarding event will therefore evoke both reinforcement-related and arousal-related responses, so their co-occurrence is expected whether or not the two are closely linked. Some of the behavioral consequences are similar. The physiological responses that follow reward may thus reflect a shared mechanism, or two independent processes triggered by a common cause. The central question, then, is not whether reinforcement and arousal are mediated by different general neuromodulatory systems, but whether the physiological consequences of reward form a tightly coupled phenomenon or can be decomposed into separable value-dependent and value-independent components.

Answering this question requires physiological markers that can separate value-dependent reinforcement processing from arousal, yet the available measures are themselves ambiguous when reward and arousal co-occur. Pupil dilation is a well-established index of LC-NE-mediated arousal (Bradley et al., 2008; McGinley et al., 2015; Joshi et al., 2016), but it also increases following reward (Bijleveld et al., 2009; Knapen et al., 2016), leaving open whether reward-evoked pupil responses reflect arousal or reward value. Cortical alpha-band activity offers complementary information, as alpha power is reliably suppressed during heightened arousal and covaries with autonomic and pupil-linked arousal (Klimesch et al., 1998; Barry et al., 2007; van Kempen et al., 2019; Compton et al., 2021). Its relationship to reward value, however, is poorly understood: reward-related oscillatory effects in human EEG have been reported most consistently in the beta band (Marco-Pallares et al., 2008; HajiHosseini & Holroyd, 2015), and whether reward value modulates alpha specifically has received little attention. A further difficulty is that most human studies use secondary rewards such as money or points, whereas animal studies typically use primary rewards such as food or water, and these engage overlapping but partly distinct circuits (Sescousse et al., 2013).

Here, we independently manipulated reward value and arousal while measuring pupil size and electroencephalographic (EEG) activity in humans. We used water delivered to the mouth as a primary reward and manipulated its value by comparing a deprivation group, in which the water was highly valued, with a no-deprivation group, in which the same water had substantially lower value. To engage arousal independently of reward value, we introduced an external arousal stimulus, the clicking sound produced by the water feeder. This design allowed us to ask whether the physiological responses elicited by reward reflect value-independent arousal, value-dependent reinforcement, or both. Because we used a primary reward, as in animal studies, it also let us examine human reward processing under conditions comparable to those used in animals.

The two measures dissociated. Reward produced pupil dilation irrespective of its value, and sound produced a comparable response, indicating that the pupillary response reflects value-independent arousal rather than reward value. EEG alpha-band activity showed a different, value-dependent pattern. Low-value reward and sound both produced widespread alpha suppression with similar temporal profiles without regional progression, consistent with a global arousal response, whereas only high-value reward increased alpha power, with activity emerging first over frontal regions and progressing posteriorly. Thus, physiological consequences of reward decompose into two dissociable components, despite their similar effects on behavior: a value-independent arousal response and a value-dependent cortical process associated with reinforcement. Contrary to the assumption that reinforcement and arousal evoked by reward are closely linked, reward engages these processes through separable mechanisms, providing a physiological framework for distinguishing their respective contributions to reward processing.

## Results

To examine the psychophysiological effects of reward and arousal, as well as their combined effect, we employed a 3 × 2 mixed design with Condition as a within-subject factor and Deprivation as a between-subject factor, all completed in a single session. The three within-subject conditions were: (1) in the water condition, only water was provided through a tube that was placed in participants’ mouths as a reward (Fig. 1); (2) in the sound condition, only a sound was presented to increase arousal levels; and (3) in the water+sound condition, both water and sound were presented simultaneously. To manipulate the reward value of water, participants were divided into two groups: the deprivation group, which was deprived of food and water for four hours prior to the experiment, and the no-deprivation group, which was not deprived. Each group completed all three stimulus conditions (water, sound, and water+sound), yielding a 3 × 2 mixed design with Condition (3 levels) as a within-subject factor and Deprivation (2 groups) as a between-subject factor, that is, six cells in total (Fig. 1B). The experiment consisted of three blocks, with each condition conducted in a separate block. During each trial, participants passively viewed a dynamic sequence of Mondrian patterns for 14 to 15 sec without any specific task (Fig. 1C). On each trial, water and/or sound was presented randomly 0 to 4 times (Fig. 1D), and changes in pupil size and EEG alpha and beta power from the pre-stimulus baseline were measured approximately 3 sec after each presentation (see Methods for details).

**Figure 1.**
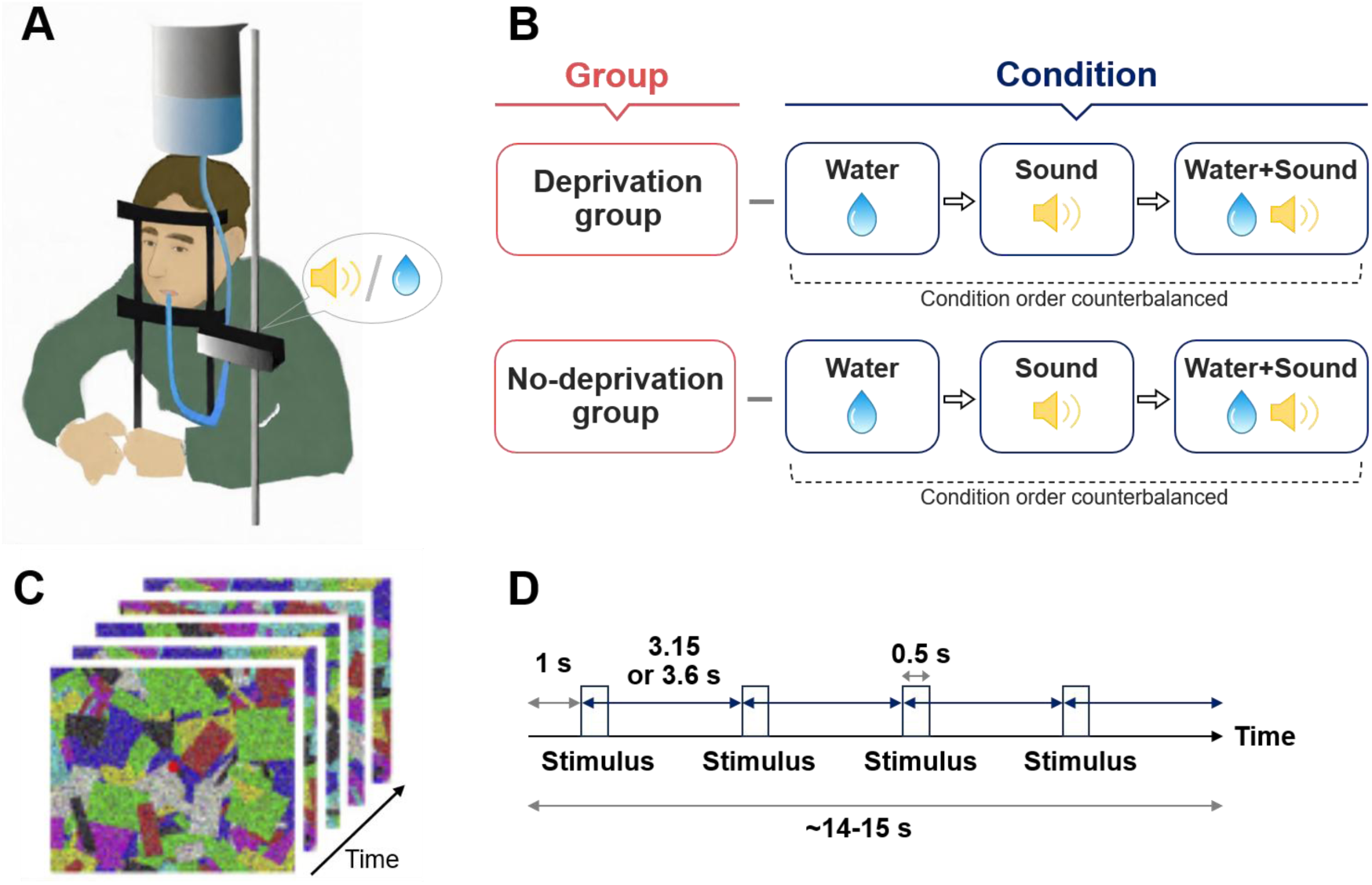
Experimental procedure. **(A)** Schematic of stimulus delivery. Water (reward) was delivered through a tube placed in the participant’s mouth. Sound (arousal) was presented as a clicking noise from the water feeder positioned behind the participant. In the water+sound condition, both stimuli were delivered simultaneously. **(B)** Experimental design. A 3 × 2 mixed design was used, with Group as a between-subject factor (deprivation and no-deprivation) and Condition as a within-subject factor (water, sound, and water+sound). The order of the three conditions was counterbalanced within each group. **(C)** Visual stimulus display. Participants fixated on a central point while passively viewing a dynamically changing background composed of Mondrian patterns with rapidly varying colored shapes. No task or response was required. **(D)** Trial structure. Each stimulus trial lasted approximately 14–15 s, during which stimuli (water and/or sound) were presented 0–4 times at predefined time points (first at 1 s after trial onset, followed by intervals of ≥3.15 s). Pupillary and EEG responses were measured relative to stimulus onset.

### Pupillary responses

Fig. 2 shows the changes in pupil size from the pre-stimulus baseline for each condition. The pupil significantly dilated in all conditions for both the deprivation and no-deprivation groups. This was revealed by one-sample t-tests at each time point followed by a cluster-based permutation test to control the familywise error across time points. The patterns of pupillary responses were very similar between the two groups. An ANOVA with Group (deprivation and no-deprivation) and Condition (water, sound, and water+sound) at each time point, followed by a cluster-based permutation test, revealed no significant clusters for the main effect of Group or the Group × Condition interaction. This indicates that deprivation had no effect on pupillary responses across conditions, and that pupillary responses do not reflect differences in reward value.

**Figure 2.**
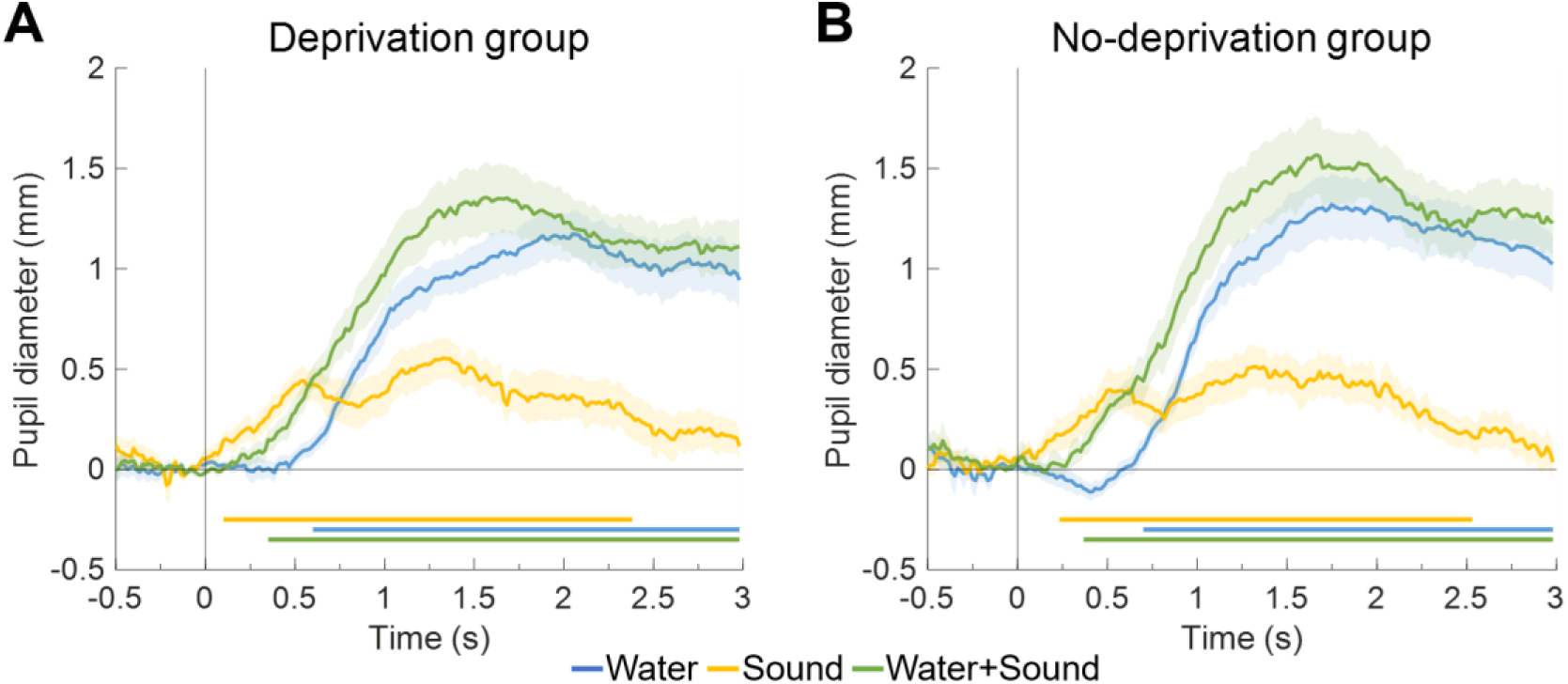
Changes in pupil size from the pre-stimulus baseline. Time courses of pupil size changes from the pre-stimulus baseline following stimulus onset, shown for water, sound, and water+sound conditions in deprivation **(A)** and no-deprivation **(B)** groups. Significant pupil dilation in all conditions across both groups, with no main effect of Group or Condition × Group interaction (ANOVA with a cluster-based permutation test). In early phases (up to around 1.25 s), pupil size increased significantly in all conditions, with no difference between groups and no interaction between condition and group. These results indicate that pupillary responses reflect value-independent arousal rather than reward value. Horizontal colored lines below plots indicate clusters of time points in which pupil size was significantly larger than zero.

The change in pupil size in the water condition was larger than in the sound condition. However, the differences in pupil size between the conditions are difficult to interpret because the water and sound stimuli differ in nature, making it challenging to match their intensities. Pupillary responses vary with stimulus intensity, such as the amount of water or the loudness and salience of the sound (Liao et al., 2016), which may have contributed to the differences observed between conditions. Thus, we did not compare or interpret the magnitude of pupil size differences across conditions (this also applies to the EEG data). Instead, we focused on the direction of the response (i.e., whether it was positive or negative) by comparing each response to zero. We observed pupil dilation in both the water and sound conditions, suggesting that pupillary responses cannot dissociate the effects of reward and arousal.

Similarly, we do not interpret the differences in the latency of pupillary responses, even though the latencies differed between conditions. This is because the time it takes for the water and sound stimuli to reach the receptors may differ. Water might take longer to reach the receptors than sound, even when both stimuli are presented simultaneously.

### EEG alpha power

We focused our primary analyses on alpha-band activity because alpha power is a well-established index of cortical state that reliably tracks arousal and attentional engagement (Klimesch et al., 1998; Barry et al., 2007; Hong et al., 2014; van Kempen et al., 2019; Compton et al., 2021; Johnston et al., 2022), while also being modulated by reward (Byrne et al., 2020; Fryer et al., 2023). This makes alpha uniquely suited for dissociating value-independent arousal effects from value-dependent reward-related processing.

Fig. 3A shows the changes in alpha power from the pre-stimulus baseline for each condition and electrode in the deprivation group. In the sound condition (shown by yellow curves in Fig. 3), alpha power generally decreased after the sound was presented. Alpha power was significantly smaller than zero in the central, parietal, and occipital channels. In contrast, in the water condition (blue curves), alpha power generally increased after the water was presented. Alpha power was significantly larger than zero in the frontal, central, and occipital channels. In the water+sound condition (green curves), alpha power appears to initially decrease after the stimuli were presented and then increased, likely reflecting a combined effect of the sound and water conditions, although the statistical tests showed no significant effect (all clusters p>.13). Together, these results indicate that, in contrast to pupillary responses, alpha power exhibits distinct patterns between the sound and water conditions: alpha power was suppressed by arousal, whereas it was enhanced by reward.

**Figure 3.**
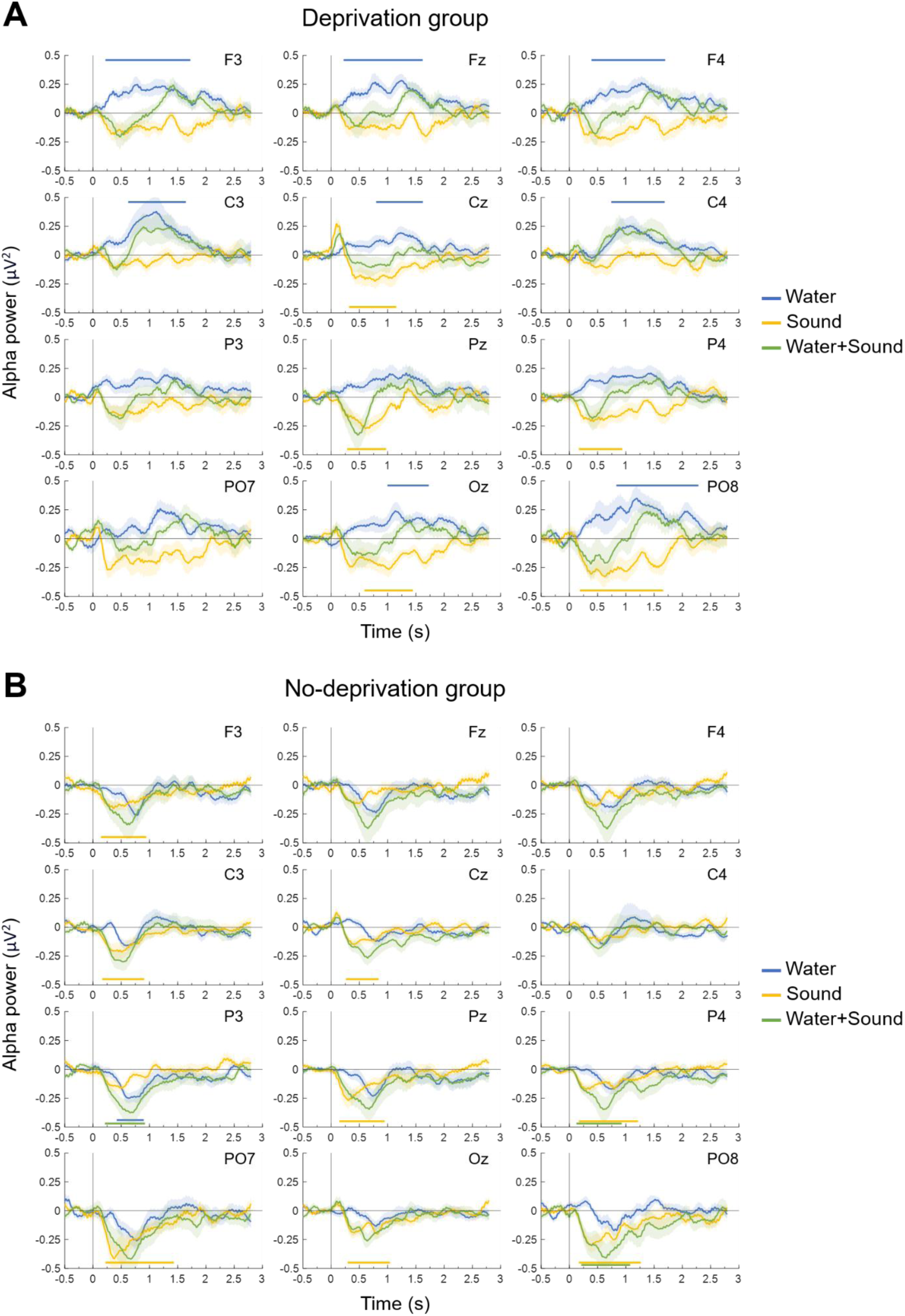
Changes in EEG alpha power from the pre-stimulus baseline. Baseline-corrected EEG alpha power (8–12 Hz) following stimulus onset, displayed across scalp electrodes (Fz, F3, F4, Cz, C3, C4, Pz, P3, P4, Oz, PO7, PO8) for each condition. **(A)** Deprivation group. Alpha power increased following water (reward) and decreased following sound (arousal), with distinct spatial distributions across electrodes. **(B)** No-deprivation group. Alpha power decreased in both water and sound conditions. Together, these results show that alpha power dissociates reward and arousal: alpha enhancement selectively reflects high-value reward whereas alpha suppression reflects arousal. Time periods during which alpha power was significantly different from zero are indicated by a blue, yellow or green horizontal line for the water, sound or water+sound condition, respectively.

However, the no-deprivation group showed a different pattern from the deprivation group (Fig. 3B). In the sound condition, alpha power decreased, similar to the deprivation group. However, in the water condition, the increase in alpha power observed in the deprivation group was not obtained; instead, alpha power generally decreased, comparable to the sound condition. A significant decrease from zero was observed at a parietal channel in the water condition. The water+sound condition also showed decreased alpha power. The similarity between the water and sound conditions in the no-deprivation group (i.e., both resulted in decreased alpha power) suggests that water may induce an arousal effect when its value is low.

Next, to examine group differences in the data shown in Fig. 3, we compared alpha power between the deprivation and no-deprivation groups for each of the sound, water and water+sound conditions (Fig. 4). In the sound (Fig. 4A) and water+sound (Fig. 4C) conditions, there were no significant differences across channels (all clusters p>.65 for the sound condition; all clusters p>.08 for the water+sound condition). In the water condition (Fig. 4B), however, we observed significant differences between the groups in several channels across the scalp. These results indicate that the increased alpha power observed in the water condition of the deprivation group reflects reward value, since this increase disappeared when water had a lower value in the no-deprivation group.

**Figure 4.**
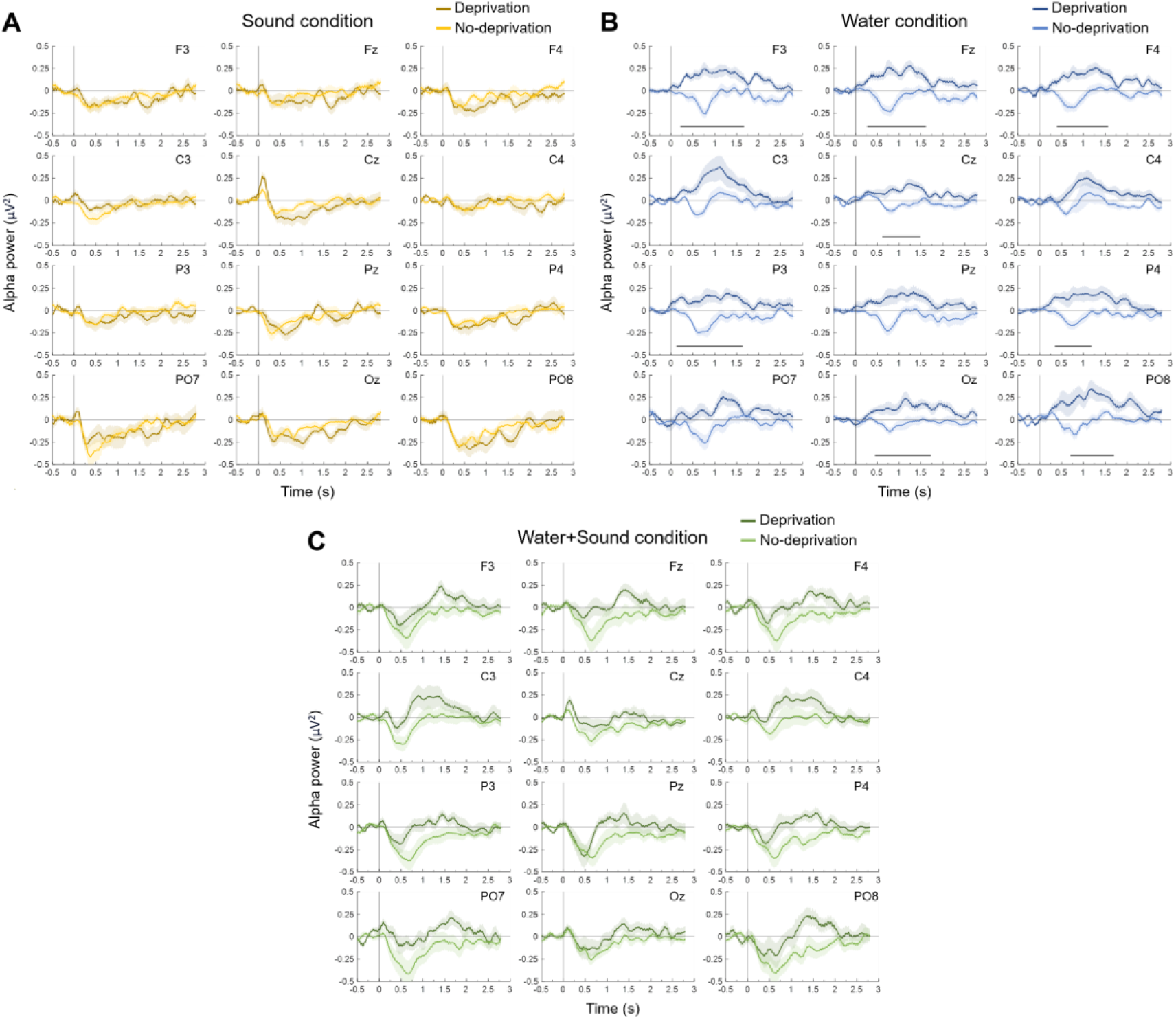
Comparison of alpha power changes between deprivation and no-deprivation groups. Differences in alpha power between groups for **(A)** sound condition, **(B)** water condition, and **(C)** water+sound condition. Time periods during which alpha power was significantly different between the two conditions are indicated as a black horizontal line. No significant group differences were observed in the sound or water+sound conditions. In contrast, the water condition showed significant differences across multiple channels, with increased alpha power present only in the deprivation group. These results indicate that alpha enhancement reflects reward value rather than arousal.

In the water condition of the deprivation group, alpha power appears to increase earlier in the frontal channels than in other regions (Fig. 3A). To investigate this further, we compared the spatiotemporal patterns of the arousal and reward effects, by examining the latency of significant clusters across electrodes (Fig. 5). In the water condition of the deprivation group, the effect initially occurred in the frontal channels, followed by the central channels, and finally the occipital channels. However, in the sound condition, the effect occurred at similar times across channels. These results suggest that the increased alpha power caused by reward initially emerges in the frontal areas and may propagate to posterior areas in a top-down manner, whereas the decreased alpha power caused by arousal emerges more simultaneously across whole areas. In the water condition of the no-deprivation group, a significant cluster was observed in only one channel, which made it infeasible to plot a latency curve. Instead, we plotted the latency of clusters with p-values less than 0.2. These clusters correspond to the largest peaks of decreased alpha power observed in each electrode. Unlike the deprivation group, the water condition in the no-deprivation group shows a flat curve, similar to the sound condition. This supports the interpretation that water induces an arousal effect in the no-deprivation group.

**Figure 5.**
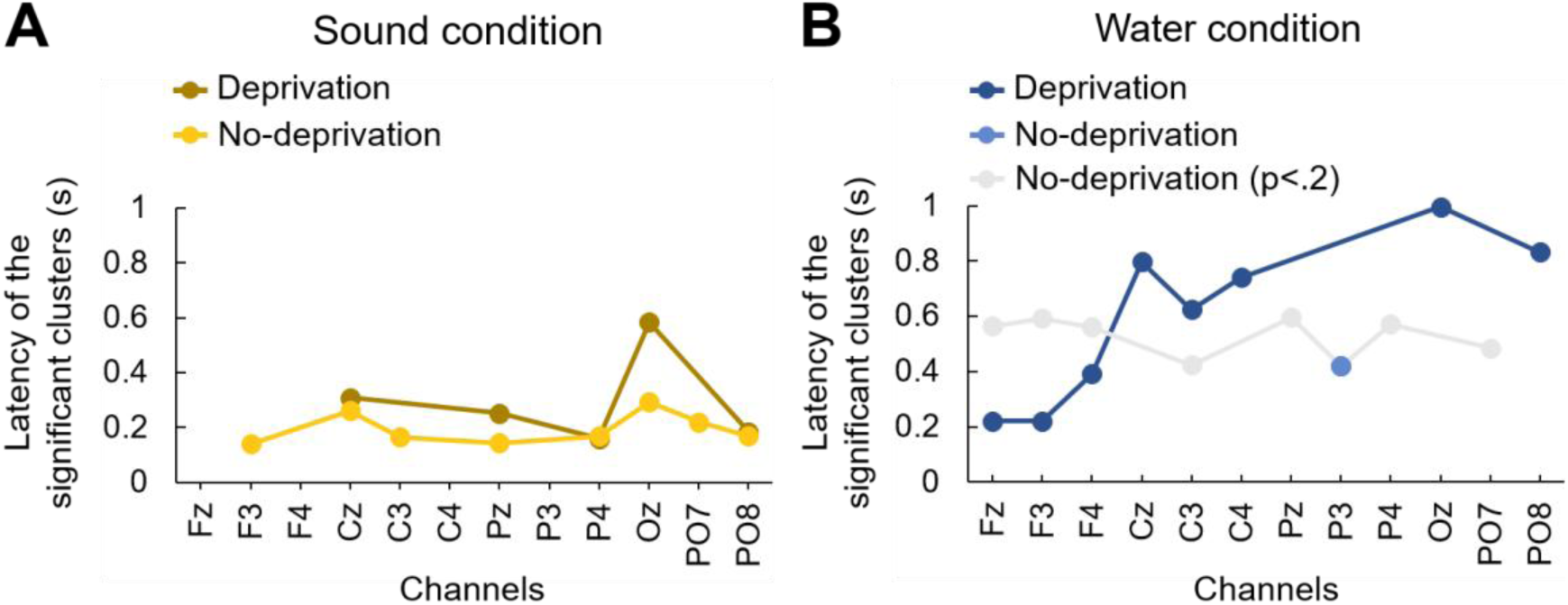
Spatiotemporal patterns of alpha power changes across electrodes. Latency of significant alpha-band modulation plotted across scalp locations from frontal to occipital electrodes in **(A)** sound condition and **(B)** water condition. In the deprivation group, alpha enhancement in the water condition emerged first in frontal electrodes and then progressed to central and occipital regions, suggesting a top-down propagation of reward-related signals. In contrast, alpha suppression in the sound condition occurred with similar timing across electrodes, consistent with a global arousal effect. The no-deprivation group showed a flat latency pattern in the water condition, resembling the sound condition.

It is possible that muscle activity associated with swallowing water introduced artifacts, which may have contributed to the increased alpha power observed in the water condition of the deprivation group. To test this possibility, we recorded EMG from the left masseter muscle and removed the corresponding ICA component in nine of twenty participants in the deprivation group. Alpha power in the water condition of the deprivation group was then compared between participants with and without EMG-based ICA removal (Fig. S1). No significant differences were observed between the two groups (all clusters p>.32), suggesting that the increased alpha power observed in the water condition of the deprivation group is not attributable to muscle artifacts.

### EEG beta power

We also examined the beta band because beta power has been linked to reward (Cohen et al., 2007; HajiHosseini & Holroyd, 2015; Marco-Pallares et al., 2008). Changes in beta power from the pre-stimulus baseline were calculated (Fig. S2). In the sound condition, no effect on beta power was observed across channels in either the deprivation or no-deprivation group (all clusters p>.40 for the deprivation group; all clusters p>.16 for the no-deprivation group). In the water and water+sound conditions, beta power increased at several channels including Oz, PO7, PO8, and C4 in the deprivation and no-deprivation groups. However, there was no difference between the deprivation and no-deprivation groups (Fig. S3; all clusters p>.37 for the water condition; all clusters p>.21 for the water+sound condition). These results indicate that beta power increased when water was provided, but this effect does not reflect reward value since it was observed in both the deprivation and no-deprivation groups. The increase in beta might instead be involved in motor processing associated with swallowing.

## Discussion

Our results demonstrate that reward-induced arousal and value-dependent reinforcement processes are largely independent, contrary to the coupling assumption, and that the effects of reward and arousal can be dissociated by alpha-band activity but not by pupillary responses. Reward delivery induced comparable pupil dilation irrespective of reward value (Fig. 2), resembling pupil dilation elicited by sound. This is the direct evidence that they reflect value-independent arousal rather than reward value. In contrast, alpha power showed a value-dependent pattern: while alpha power decreased in response to sound and low-value reward, it increased selectively when reward value was high, namely in the deprivation condition (Figs. 3 and 4). Furthermore, the distinct spatiotemporal patterns of alpha modulation (Fig. 5) suggest that these effects arise from different neural mechanisms. Alpha suppression occurred nearly simultaneously across widespread cortical regions, consistent with global arousal signals (Reimer et al., 2014; McGinley et al., 2015), whereas alpha enhancement emerged first in frontal regions and then propagated posteriorly, suggesting a top-down process related to reward value, potentially mediated by prefrontal control over sensory cortices (Serences, 2008). Together, these findings indicate that reward induces both value-independent arousal resembling that elicited by sound and value-dependent cortical processing through independent mechanisms, providing a physiological framework for understanding how these processes coexist without being closely coupled.

The present findings clarify the relationship between arousal-related and reward-value-related signals in the frameworks of cortical processing. While prior work established that both processes are co-triggered by reward, whether they are closely coupled or operate independently could not be resolved from paradigms using reward alone. Previous work has suggested that arousal-related signals, linked to motivational significance, can modulate cortical gain and sensory processing and are reflected in pupil dilation (e.g., Nassar et al., 2012). The current results extend this perspective by showing that such value-independent arousal signals do not fully account for reward-value-related processing. While pupil-linked responses captured arousal effects across conditions, alpha-band dynamics revealed an additional value-dependent cortical process, indicating that reward engages mechanisms beyond those indexed by arousal alone.

These findings also speak to broader question of how reward signals are organized in the brain. Prior work has shown that reward enhances perceptual learning and plasticity, particularly under conditions of increased motivational value (Seitz et al., 2009; Law & Gold, 2009; Shibata et al., 2011; Kahnt et al., 2011; Herpers et al., 2021; Murris et al., 2021; Kim et al., 2026). The present dissociation suggests that two distinct processes may contribute to such effects. Arousal-related mechanisms may facilitate these processes by increasing global gain or sensitivity (Aston-Jones & Cohen, 2005; Nassar et al., 2012), whereas value-dependent processes may provide an additional enhancement when reward is highly valued. This common-input, independent-outputs account may help reconcile differences between theories emphasizing reinforcement signals (Bromberg-Martin et al., 2010; Wise, 2004; Watabe-Uchida et al., 2017; Seitz et al., 2009; Law & Gold, 2009; Herpers et al., 2021; Murris et al., 2021; Kim et al., 2026) and those emphasizing arousal or attention (Aston-Jones & Cohen, 2005; Nassar et al., 2012).

The present results also help explain inconsistencies in prior studies. Some studies have reported value-dependent modulation of pupil size, particularly when reward magnitude is unpredictable or embedded in oddball paradigms (Knapen et al., 2016; Cole et al., 2022). Under such conditions, pupil responses may reflect variations in surprise or attentional allocation rather than reward value per se (Preuschoff et al., 2011). In contrast, the present design minimized unpredictability about stimulus values, allowing a clearer isolation of arousal effects. Similarly, studies using monetary rewards have often reported alpha suppression rather than enhancement (Byrne et al., 2020; Fryer et al., 2023). One possible explanation is that the effective value of such rewards is relatively low, making their physiological effects resemble arousal. Another possibility is that primary and secondary rewards engage partially distinct neural mechanisms. While both involve overlapping reward-related regions such as the ventral tegmental area, ventral striatum (Schultz et al., 1997; Haber & Knutson, 2010), and insula (Craig, 2009), secondary rewards rely more strongly on orbitofrontal cortex representations (Sescousse et al., 2013; Wallis, 2007).

Beta-band activity has also been associated with reward processing. Beta power increases in response to monetary reward cues (Cohen et al., 2007; HajiHosseini & Holroyd, 2015; Marco-Pallares et al., 2008). The present results found no clear value-dependent beta modulation. Beta power did not change following sound but increased following water delivery irrespective of deprivation state, raising the possibility that the beta response may reflect motor-related cortical activity or muscle artifacts associated with swallowing rather than reward value.

At the neural level, the dissociation observed here is consistent with the organization of arousal and reward systems. The global and temporally uniform alpha suppression associated with arousal is consistent with the locus coeruleus–norepinephrine system, which projects diffusely across the cortex and modulates neural activity on a rapid timescale (Aston-Jones & Cohen, 2005; Berridge & Waterhouse, 2003). In contrast, the anterior-to-posterior progression of alpha enhancement is consistent with reward-related signaling pathways in which midbrain dopaminergic projections from the ventral tegmental area reach prefrontal cortex, which in turn exerts top-down influences on sensory cortices via cortico-cortical feedback pathways (Schultz et al., 1997; Noudoost & Moore, 2011). However, recent work suggests that dopamine may encode broader variables such as salience or motivational relevance rather than reward value alone (Bromberg-Martin et al., 2010; Gershman & Uchida, 2019; Gershman et al., 2024). Thus, while the present findings are consistent with the involvement of reward-related neuromodulatory systems, the precise neurochemical mechanisms underlying alpha enhancement remain to be determined.

The functional role of reward-related alpha enhancement remains an important question. Alpha oscillations have been associated with inhibitory processing and reduced neuronal firing in cortical circuits (Klimesch et al., 2007; Jensen & Mazaheri, 2010; van Kerkoerle et al., 2014). From this perspective, increased alpha power may reflect selective suppression of irrelevant or competing information, thereby enhancing the signal-to-noise ratio of sensory processing. Consistent with this interpretation, reward has been shown to modulate sensory cortical activity and improve perceptual processing (Serences, 2008; Seitz et al., 2009; Law & Gold, 2009; Kahnt et al., 2011). Thus, value-dependent alpha enhancement may reflect a mechanism through which reward selectively shapes cortical processing beyond global arousal effects.

Several limitations should be noted. First, EEG does not provide direct access to underlying neuromodulatory systems, making it difficult to determine the precise neurochemical basis of the observed effects. Second, although the dissociation between arousal and reward-related processes is clear at the physiological level, the functional role of alpha enhancement in cognition and behavior requires further investigation. Third, the observed frontal-to-posterior latency progression in alpha enhancement (Fig. 5) is consistent with top-down propagation but does not establish it: latency ordering across scalp electrodes is correlational and cannot rule out alternative explanations such as regionally distinct intrinsic time constants. Fourth, reward value was manipulated categorically (deprivation vs. no-deprivation) rather than parametrically, so the term "value-dependent" reflects a two-level contrast rather than a graded value function, and claims should be read accordingly.

In summary, the present study dissociates two fundamental components of reward processing: a value-independent arousal response resembling that elicited by sound and a value-dependent cortical process. While arousal-related signals are reflected in pupil dilation and global alpha suppression, reward value is selectively reflected in alpha enhancement with a distinct spatiotemporal profile. This dissociation provides a unified physiological framework for understanding how reward-related processes coexist in the brain, and suggests that reward influences neural processing through multiple, functionally distinct mechanisms. Importantly, the distinctness of these two processes alleviates concerns that reward-based studies of reinforcement learning are confounded by arousal: the value-dependent signal identified here reflects value-dependent reinforcement processes proper, not a byproduct of generalized arousal.

## Materials and Methods

### Participants

A total of 52 subjects participated in the study. In the deprivation group, 12 subjects (mean age 25.1 years, age range 18–48 years, 4 females) participated in the pupil experiment, and 20 subjects (mean age 23.4 years, age range 18–46 years, 14 females) participated in the EEG experiment. In the no-deprivation group, 20 subjects (mean age 26.4 years, age range 18–50 years, 13 females) participated in the experiment. All participants had normal or corrected-to-normal vision, and were provided written informed consent. The study was approved by the institutional review board at Brown University.

### Stimuli and apparatus

Water was delivered to participants’ mouths through a tube connected to a water feeder (Fig. 1A). The delivery of water was controlled using ValveLink8.2 system (Automate Scientific, Inc.). Clicking sounds were produced by the water feeder when its gate opened and closed. Visual stimuli were presented on an LCD monitor (resolution: 1920 x 1080; refresh rate: 60 Hz) with a gray background at a viewing distance of 57 cm. Participants put their heads on a chin rest during experiments. The experiments were conducted in a shielded and soundproof room.

### Procedures

All participants underwent the following three conditions. In the water condition, approximately 1 ml of water was provided over 500 ms. The water feeder was placed outside the room to prevent participants from hearing the clicking sound. Participants were instructed to swallow the water when it was provided. In the sound condition, a clicking sound from the water feeder, which was positioned directly behind participants, was presented to increase arousal levels. A tube was placed in participants’ mouths to simulate the water condition, but the water feeder was empty and no water was provided. The sound occurred twice during each presentation— once when the feeder’s gate opened and once when it closed—with a 500 ms interval between the two sounds. In the water+sound condition, the water and sound were presented simultaneously. The water feeder was positioned directly behind participants like the sound condition. To manipulate reward value levels of water, participants were divided into two groups: those in the deprivation group were deprived of food and water for four hours prior to the experiment, while those in the no-deprivation group were not. Both groups completed all three stimulus conditions, resulting in a 3 × 2 mixed design with Condition as a within-subject factor and Deprivation as a between-subject factor (Fig. 1B).

During each trial, participants passively viewed a dynamic sequence of Mondrian patterns (Fig. 1C). Each Mondrian pattern consisted of 300 randomly placed, physically overlapping rectangles or ellipses of various sizes, presented at a rate of 20 Hz. Additionally, colored pixel noise covered 50% of the entire screen. The Mondrian patterns were used because we were interested in understanding the effect of reward and arousal on visual processing, and the same stimulus was used in a previous study examining the effect of reward on visual perceptual learning (Seitz et al., 2009). Participants were instructed to fixate on a point that appeared at the center of the screen during the trial.

There were two types of trials: stimulus trials and no-stimulus trials, which were presented alternately in each block. In a stimulus trial, water and/or sound was presented randomly from 0 to 4 times (Fig. 1D). Each trial contained four possible time points at which a stimulus (i.e., water and/or sound) could occur. The first time point was 1 sec after trial onset, and each of the remaining three occurred after a random interval of either 3.15 or 3.6 sec from the previous time point. Thus, the interval between stimuli was at least 3.15 sec (in cases where stimuli were presented consecutively). Each stimulus trial lasted approximately 14 to 15 sec, during which pupillary and EEG responses to water and/or sound were recorded. No-stimulus trials were the same as stimulus trials except that neither water nor sound was presented and their duration was 7.3 sec. No-stimulus trials were included to replicate the design of the previous study (Seitz et al., 2009).

The three conditions (i.e., water, sound, and water+sound) were conducted in three separate blocks. Each block consisted of 27 repetitions of alternating stimulus and no-stimulus trials. Participants were allowed to take breaks between trials as needed and also took a 3-min break between blocks. The total number of stimulus presentations (i.e., water and/or sound) for each condition was 69. The order of the conditions was approximately counterbalanced within each of the deprivation and no-deprivation groups.

In the deprivation group, the experiments measuring pupillary responses and EEG were conducted separately since we could not record them simultaneously due to the limitations of the devices. The procedures for the two experiments were the same, except for whether pupil size or EEG was measured. In the no-deprivation group, both pupil size and EEG were recorded in the same experiment.

### Pupil recording and analysis

Pupil data were recorded using a Tobii Pro Nano (Tobii). Pupil diameter of both eyes was recorded at 60 Hz, and the data from both eyes were averaged. Missing data caused by blinks were linearly interpolated from 50 ms before the starting point to 50 ms after the endpoint of the blink. Trials with missing data exceeding 167 ms were excluded from the analysis. Pupil size data were segmented for each trial from 500 ms before stimulus (water or sound) onset to 3 sec after stimulus onset. The average pupil size during the pre-stimulus baseline (-167 ms to 0 ms) was subtracted from all time points on each trial. The data were then averaged across trials for each condition.

### EEG recording and analysis

EEG data were recorded using an ActiCap electrodes system and a BrainAmp amplifier (Brain Products GmbH). The following 12 sites from the 10/10 system were used: Fz, F3, F4, Cz, C3, C4, Pz, P3, P4, Oz, PO7, and PO8. CPz was recorded as reference. EOG and EMG data were recorded using bipolar passive electrodes and a BrainAmp ExG amplifier (Brain Products GmbH). Horizontal EOG was recorded from electrodes placed lateral to the external canthi to detect horizontal eye movements, and vertical EOG was recorded from electrodes placed below and above the left eye to detect blinks and vertical eye movements. EMG was recorded from electrodes placed on the left masseter muscle to detect muscle activity during water swallowing. EMG from the masseter was recorded for nine out of twenty participants in the deprivation group and all participants in the no-deprivation group. The signals were sampled at a rate of 1,000 Hz and filtered online between 0.03 and 100 Hz.

Preprocessing was performed using the EEGLAB Toolbox (Delorme & Makeig, 2004). All the signals were bandpass filtered between 1 and 30 Hz and resampled at 250 Hz. Noisy channels were removed by visual inspection and were interpolated by the mean signals of their neighboring channels (a single channel was removed from one participant in the deprivation group; none were removed in the no-deprivation group). EEG data were segmented in epochs from 1 sec before the stimulus onset to 3 sec after the stimulus onset. The EEG data was re-referenced off-line to the average of all scalp electrodes. Independent component analysis (ICA) was performed to remove components involved in blinks, eye movements, and water swallowing.

Alpha and beta power were calculated using temporal spectral evolution (TSE; Salmelin & Hari, 1994). The preprocessed EEG data for each trial were bandpass filtered between 8 and 12 Hz (alpha) or 20 and 30 Hz (beta). The data were further segmented into epochs from -500 ms to 2.8 sec relative to the stimulus onset to remove filter warm-up artifacts at the edges of the epoch window. The data were then rectified and smoothed with moving average of a 100-ms window. The average power of the pre-stimulus baseline (-500 ms to 0 ms) was subtracted from all time points for each trial. The resulting data were averaged across trials for each individual participant.

### Statistical analysis

We used a mass univariate analysis with a non-parametric cluster-based permutation test (Maris & Oostenveld, 2007; Groppe et al., 2011) to statistically test the time-series data of both pupil size and EEG. This approach controls the familywise error across time points and electrodes. To compare changes in pupil size and EEG alpha and beta power against the chance level (0), we conducted one-sample, two-tailed t tests at each time point (from stimulus onset to the end of the epoch) for each condition. Clusters were formed by grouping consecutive time points where the single-point t tests were significant (p<.05), and the t scores within each cluster were summed to obtain a cluster-level t mass. Each cluster-level t mass was then compared with a null distribution of cluster-level t mass, which was estimated using the maximum cluster-level t mass calculated in each of 2500 random permutations of the original data. The p value of each cluster was calculated based on its rank in the null distribution, and clusters with p value less than 0.05 were reported as significant.

For pupil size, to compare the results between the deprivation and no-deprivation groups, we conducted an ANOVA at each time point and applied a cluster-based permutation test (Fields & Kuperberg, 2019). The procedure for the cluster-based permutation test was the same as that used in t tests, except that F values were used instead of t values. For alpha and beta power, two-sample, two-tailed t tests were conducted with a cluster-based permutation test for each condition to compare the results between the deprivation and no-deprivation groups.

## Acknowledgments

This study was supported by NIH R01EY019466 (to T.W.), R01EY027841 (to T.W.), R01EY031705 (to Y.S.), and NSF-BSF BCS2241417 (to T.W.).

## Supplemental Material

**Figure S1.**
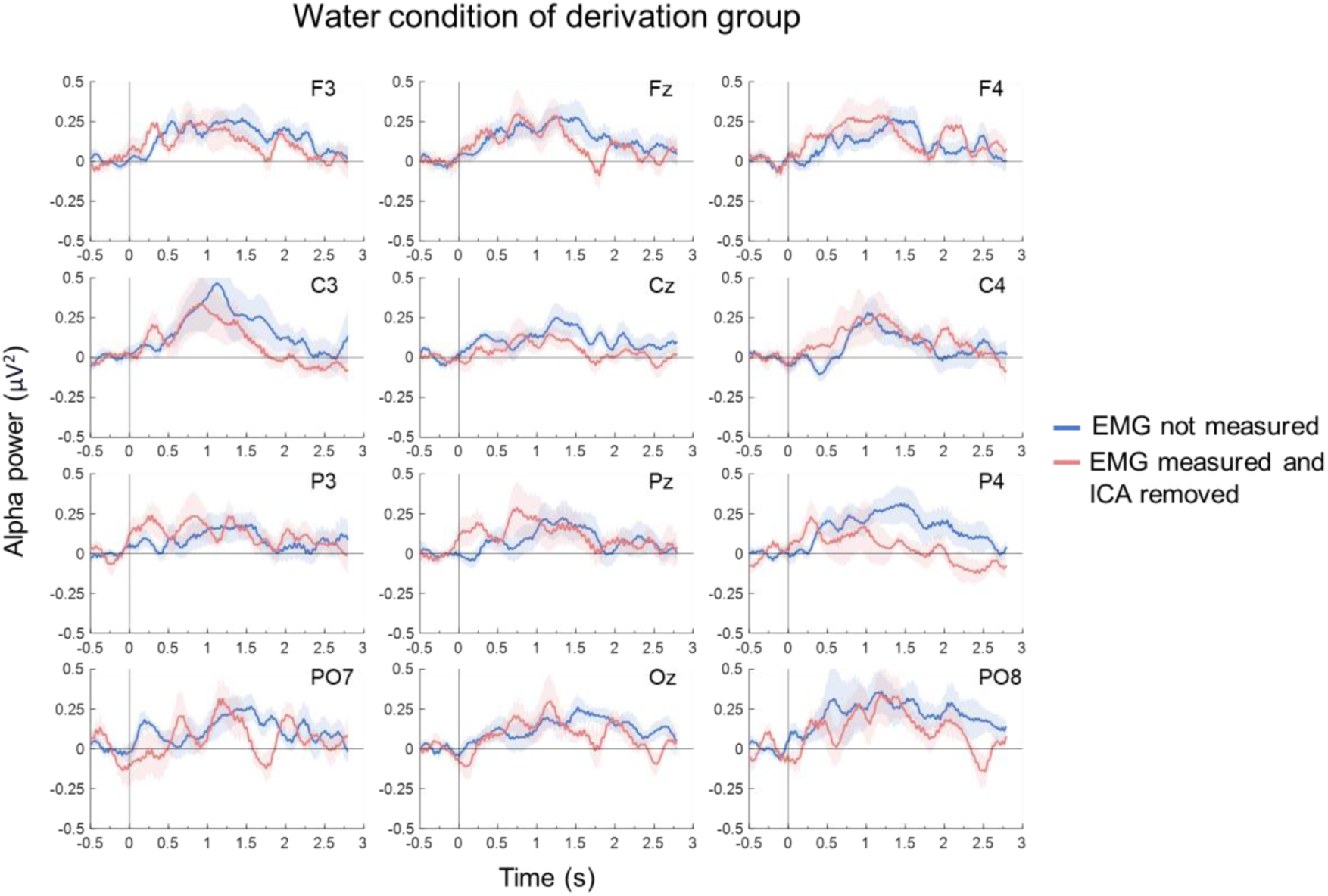
Control analysis of EMG-related artifacts in the water condition. Comparison of alpha power changes in the deprivation group (water condition) between participants with and without EMG-based ICA removal of masseter activity. No significant differences were observed, indicating that increased alpha power in the water condition is not attributable to muscle artifacts associated with swallowing.

**Figure S2.**
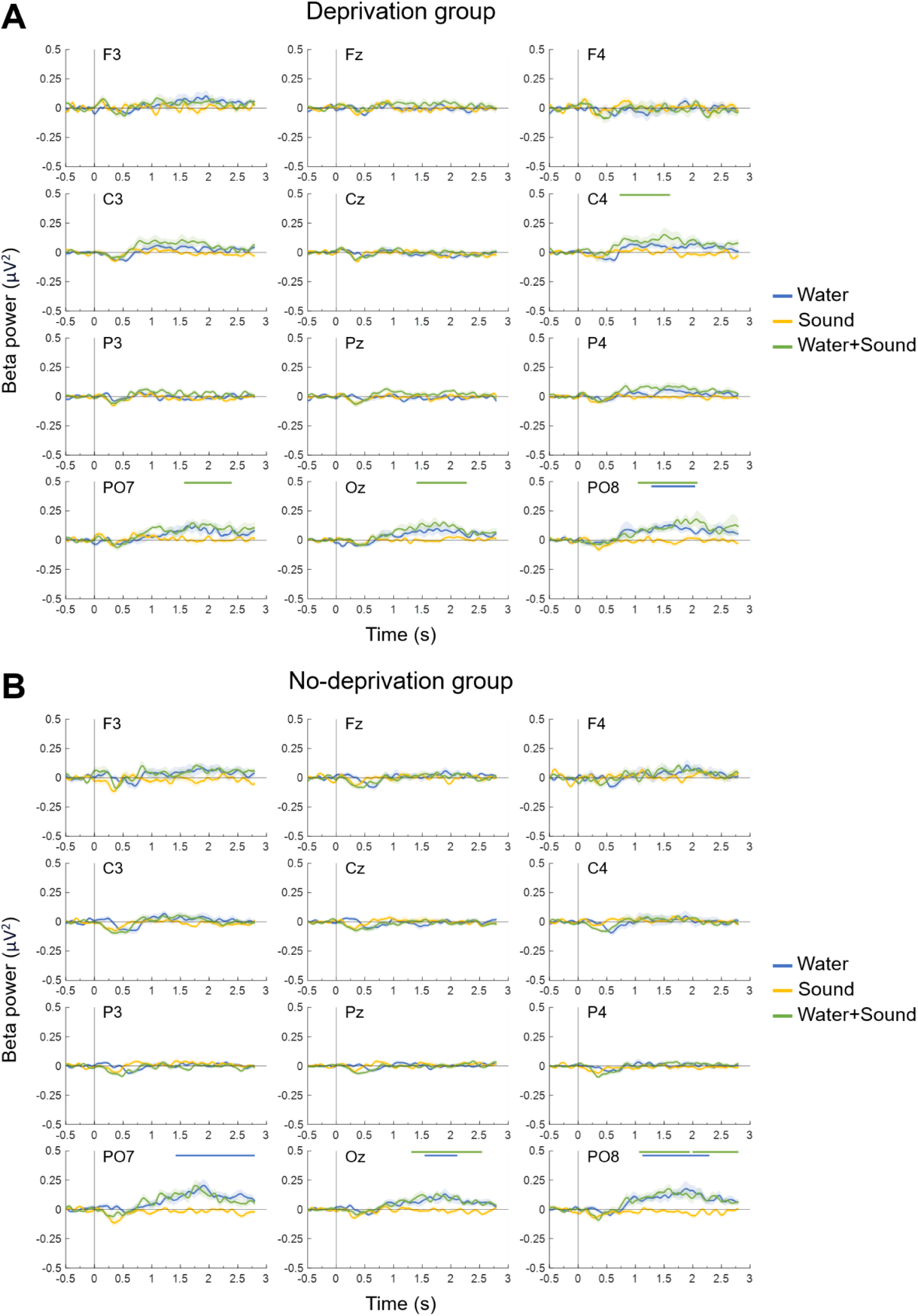
The changes in beta power from the pre-stimulus baseline. Baseline-corrected EEG beta power (20–30 Hz) following stimulus onset, shown separately for the water (reward), sound (arousal), and water+sound conditions across scalp electrodes (Fz, F3, F4, Cz, C3, C4, Pz, P3, P4, Oz, PO7, PO8) in **(A)** deprivation and **(B)** no-deprivation groups. In the sound condition, no significant modulation across electrodes in either group (all clusters p>.40 in the deprivation group; p>.16 in the no-deprivation group). In the water condition, increases in beta power were observed at occipital electrodes (e.g., Oz, PO7, PO8) in both groups. In the water+sound condition, similar increases in beta power were observed across overlapping electrode sites in both groups. No group differences were observed across all conditions, indicating that beta power changes were independent of reward value, with beta increases likely reflecting motor-related processes such as swallowing.

**Figure S3.**
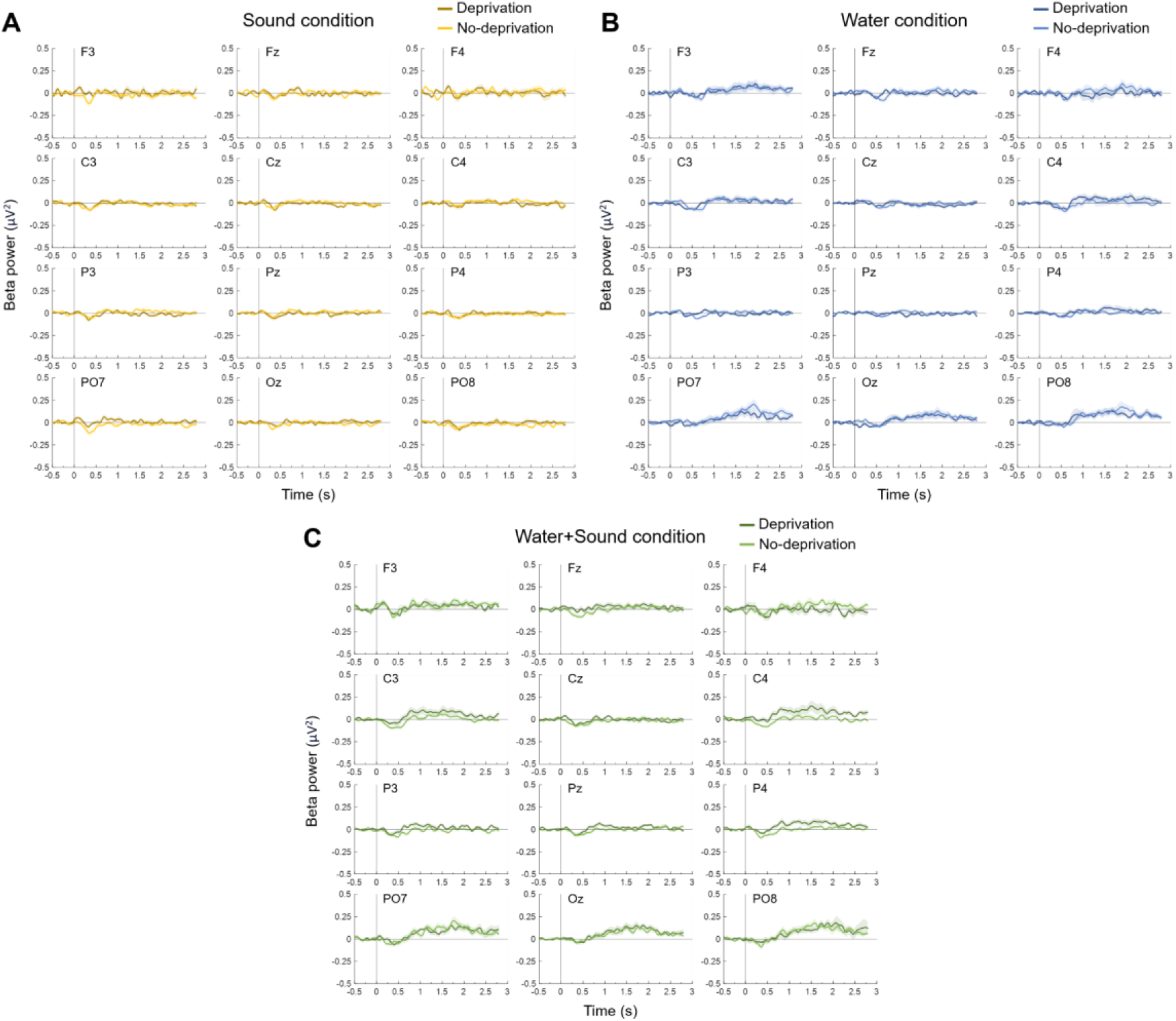
Comparison of beta power changes between deprivation and no-deprivation groups. Differences in baseline-corrected EEG beta power (20–30 Hz) between deprivation and no-deprivation groups, shown for the **(A)** sound, **(B)** water (reward) and **(C)** water+sound conditions across scalp electrodes. No significant differences in beta power were observed between groups for any of the three conditions. This confirms that beta modulation does not reflect reward value.

## References

1. Aston-Jones G, Cohen JD (2005) An integrative theory of locus coeruleus–norepinephrine function: Adaptive gain and optimal performance. Annu Rev Neurosci 28:403–450.

2. Barry RJ, Clarke AR, Johnstone SJ, Magee CA, Rushby JA (2007) EEG differences between eyes-closed and eyes-open resting conditions. Clin Neurophysiol 118:2765–2773.

3. Berridge CW, Schmeichel BE, España RA (2012) Noradrenergic modulation of wakefulness/arousal. Sleep Med Rev 16:187–197.

4. Berridge CW, Waterhouse BD (2003) The locus coeruleus–noradrenergic system: Modulation of behavioral state and state-dependent cognitive processes. Brain Res Rev 42:33–84.

5. Bijleveld E, Custers R, Aarts H (2009) The unconscious eye opener: pupil dilation reveals strategic recruitment of resources upon presentation of subliminal reward cues. Psychol Sci 20:1313–1315.

6. Bradley MM, Miccoli L, Escrig MA, Lang PJ (2008) The pupil as a measure of emotional arousal and autonomic activation. Psychophysiol 45:602–607.

7. Breton-Provencher V, Drummond GT, Feng J, Li Y, Sur M (2022) Spatiotemporal dynamics of noradrenaline during learned behaviour. Nature 606:732–738.

8. Bromberg-Martin ES, Matsumoto M, Hikosaka O (2010) Dopamine in motivational control: Rewarding, aversive, and alerting. Neuron 68:815–834.

9. Byrne A, Kokmotou K, Roberts H, Soto V, Tyson-Carr J, Hewitt D, Giesbrecht T, Stancak A (2020) The cortical oscillatory patterns associated with varying levels of reward during an effortful vigilance task. Exp Brain Res 238:1839–1859.

10. Carter ME, Yizhar O, Chikahisa S, Nguyen H, Adamantidis A, Nishino S, Deisseroth K, de Lecea L (2010) Tuning arousal with optogenetic modulation of locus coeruleus neurons. Nat Neurosci 13:1526–1533.

11. Cohen MX, Elger CE, Ranganath C (2007) Reward expectation modulates feedback-related negativity and EEG spectra. NeuroImage 35:968–978.

12. Cole L, Lightman S, Clark R, Gilchrist ID (2022) Tonic and phasic effects of reward on the pupil: implications for locus coeruleus function. Proc Biol Sci 289:20221545.

13. Compton RJ, Gearinger D, Wild H, Rette D, Heaton EC, Histon S, Thiel P, Jaskir M (2021) Simultaneous EEG and pupillary evidence for post-error arousal during a speeded performance task. Eur J Neurosci 53:543–555.

14. Craig AD (2009) How do you feel—now? The anterior insula and human awareness. Nat Rev Neurosci 10:59–70.

15. Critchley HD, Mathias CJ, Dolan RJ (2001) Neural activity in the human brain relating to uncertainty and arousal during anticipation. Neuron 29:537–545.

16. Delorme A, Makeig S (2004) EEGLAB: An open source toolbox for analysis of single-trial EEG dynamics including independent component analysis. J Neurosci Methods 134:9–21.

17. Fields EC, Kuperberg GR (2019) Having your cake and eating it too: flexibility and power with mass univariate statistics for ERP data. Psychophysiology 57:e13468.

18. Fryer SL, Marton TF, Roach BJ, Holroyd CB, Abram SV, Lau KJ, Ford JM, McQuaid JR, Mathalon DH (2023) Alpha event-related desynchronization during reward processing in schizophrenia. Biol Psychiatry Cogn Neurosci Neuroimaging 8:551–559.

19. Gershman SJ, Uchida N (2019) Believing in dopamine. Nat Rev Neurosci 20:703–714.

20. Gershman SJ, Assad JA, Datta SR, Linderman SW, Sabatini BL, Uchida N, Wilbrecht L (2024) Explaining dopamine through prediction errors and beyond. Nat Neurosci 27:1645– 1655.

21. Glennon E, Carcea I, Martins ARO, Multani J, Shehu I, Svirsky MA, Froemke RC (2019) Locus coeruleus activation accelerates perceptual learning. Brain Res 1709:39–49.

22. Groppe DM, Urbach TP, Kutas M (2011) Mass univariate analysis of event-related brain potentials/fields I: a critical tutorial review. Psychophysiology 48:1711–1725.

23. Haber SN, Knutson B (2010) The reward circuit: Linking primate anatomy and human imaging. Neuropsychopharmacology 35:4–26.

24. HajiHosseini A, Holroyd CB (2015) Reward feedback stimuli elicit high-beta EEG oscillations in human dorsolateral prefrontal cortex. Sci Rep 5:13021.

25. Herpers J, Arsenault JT, Vanduffel W, Vogels R (2021) Stimulation of the ventral tegmental area induces visual cortical plasticity at the neuronal level. Cell Rep 37:109998.

26. Hong L, Walz JM, Sajda P (2014) Your eyes give you away: prestimulus changes in pupil diameter correlate with poststimulus task-related EEG dynamics. PLoS One 9:e91321.

27. Jensen O, Mazaheri A (2010) Shaping functional architecture by oscillatory alpha activity: gating by inhibition. Front Hum Neurosci 4:186.

28. Johnston R, Snyder AC, Schibler RS, Smith MA (2022) EEG signals index a global signature of arousal embedded in neuronal population recordings. eNeuro 9:ENEURO.0012-22.2022.

29. Joshi S, Li Y, Kalwani RM, Gold JI (2016) Relationships between pupil diameter and neuronal activity in the locus coeruleus, colliculi, and cingulate cortex. Neuron 89:221–234.

30. Kahnt T, Grueschow M, Speck O, Haynes JD (2011) Perceptual learning and decision-making in human medial frontal cortex. Neuron 70:549–559.

31. Kim D, Wang Z, Sakagami M, Sasaki Y, Watanabe T (2026) Motivational state determines error-sensitive learning modes in visual perceptual learning. Cereb Cortex 36:bhag006.

32. Klimesch W, Doppelmayr M, Russegger H, Pachinger T, Schwaiger J (1998) Induced alpha band power changes in the human EEG and attention. Neurosci Lett 244:73–76.

33. Klimesch W, Sauseng P, Hanslmayr S (2007) EEG alpha oscillations: the inhibition–timing hypothesis. Brain Res Rev 53:63–88.

34. Knapen T, de Gee JW, Brascamp J, Nuiten S, Hoppenbrouwers S, Theeuwes J (2016) Cognitive and ocular factors jointly determine pupil responses under equiluminance. PLoS One 11:e0155574.

35. Law CT, Gold JI (2009) Reinforcement learning can account for associative and perceptual learning on a visual-decision task. Nat Neurosci 12:655–663.

36. Liao HI, Kidani S, Yoneya M, Kashino M, Furukawa S (2016) Correspondences among pupillary dilation response, subjective salience of sounds, and loudness. Psychon Bull Rev 23:412–425.

37. Marco-Pallares J, Cucurell D, Cunillera T, García R, Andrés-Pueyo A, Münte TF, Rodríguez-Fornells A (2008) Human oscillatory activity associated to reward processing in a gambling task. Neuropsychologia 46:241–248.

38. Maris E, Oostenveld R (2007) Nonparametric statistical testing of EEG-and MEG-data. J Neurosci Methods 164:177–190.

39. Mather M, Sutherland MR (2011) Arousal-biased competition in perception and memory. Perspect Psychol Sci 6:114–133.

40. Matsumoto M, Hikosaka O (2009) Two types of dopamine neuron distinctly convey positive and negative motivational signals. Nature 459:837–841.

41. McGinley MJ, Vinck M, Reimer J, Batista-Brito R, Zagha E, Cadwell CR, Tolias AS, Cardin JA, McCormick DA (2015) Waking state: rapid variations modulate neural and behavioral responses. Neuron 87:1143–1161.

42. Murris SR, Arsenault JT, Raman R, Vogels R, Vanduffel W (2021) Electrical stimulation of the macaque ventral tegmental area drives category-selective learning without attention. Neuron 109:1381–1395.

43. Nassar MR, Rumsey KM, Wilson RC, Parikh K, Heasly B, Gold JI (2012) Rational regulation of learning dynamics by pupil-linked arousal systems. Nat Neurosci 15:1040–1046.

44. Noudoost B, Moore T (2011) Control of visual cortical signals by prefrontal dopamine. Nature 474:372–375.

45. Preuschoff K, ’t Hart BM, Einhäuser W (2011) Pupil dilation signals surprise: evidence for noradrenaline’s role in decision making. Front Neurosci 69:182–193.

46. Reimer J, Froudarakis E, Cadwell CR, Yatsenko D, Denfield GH, Tolias AS (2014) Pupil fluctuations track fast switching of cortical states during quiet wakefulness. Neuron 84:355– 362.

47. Salmelin R, Hari R (1994) Spatiotemporal characteristics of sensorimotor neuromagnetic rhythms related to thumb movement. Neuroscience 60:537–550.

48. Sara SJ (2009) The locus coeruleus and noradrenergic modulation of cognition. Nat Rev Neurosci 10:211–223.

49. Schultz W, Dayan P, Montague PR (1997) A neural substrate of prediction and reward. Science 275:1593–1599.

50. Schultz W (2016) Dopamine reward prediction-error signalling: a two-component response. Nat Rev Neurosci 17:183–195.

51. Seitz AR, Kim D, Watanabe T (2009) Rewards evoke learning of unconsciously processed visual stimuli in adult humans. Neuron 61:700–707.

52. Serences JT (2008) Value-based modulations in human visual cortex. Neuron 60:1169– 1181.

53. Sescousse G, Caldú X, Segura B, Dreher JC (2013) Processing of primary and secondary rewards: a quantitative meta-analysis and review of human functional neuroimaging studies. Neurosci Biobehav Rev 37:681–696.

54. Shibata K, Watanabe T, Sasaki Y, Kawato M (2011) Perceptual learning incepted by decoded fMRI neurofeedback without stimulus presentation. Science 334:1413–1415.

55. van Kempen J, Loughnane GM, Newman DP, Kelly SP, Thiele A, O’Connell RG, Bellgrove MA (2019) Behavioural and neural signatures of perceptual decision-making are modulated by pupil-linked arousal. Elife 8:e42541.

56. van Kerkoerle T, Self MW, Dagnino B, Gariel-Mathis MA, Poort J, van der Togt C, Roelfsema PR (2014) Alpha and gamma oscillations characterize feedback and feedforward processing in monkey visual cortex. Proc Natl Acad Sci USA 111:14332–14341.

57. Wallis JD (2007) Orbitofrontal cortex and decision making. Annu Rev Neurosci 30:31–56.

58. Watabe-Uchida M, Eshel N, Uchida N (2017) Neural circuitry of reward prediction error. Annu Rev Neurosci 40:373–394.

59. Wise RA (2004) Dopamine, learning, and motivation. Nat Rev Neurosci 5:483–494.

